# Investigating causal relationships between sleep traits and risk of breast cancer: a Mendelian randomization study

**DOI:** 10.1101/457572

**Authors:** Rebecca C. Richmond, Emma L. Anderson, Hassan S. Dashti, Samuel E. Jones, Jacqueline M. Lane, Linn Beate Strand, Ben Brumpton, Martin Rutter, Andrew R. Wood, Caroline L. Relton, Marcus Munafò, Timothy M. Frayling, Richard M. Martin, Richa Saxena, Michael N. Weedon, Debbie A. Lawlor, George Davey Smith

## Abstract

**Objective:** To examine whether sleep traits have a causal effect on risk of breast cancer.

**Design:** Multivariable regression, one- and two-sample Mendelian randomization.

**Setting:** The UK Biobank prospective cohort study and the Breast Cancer Association Consortium (BCAC) case-control genome-wide association study.

**Participants:** 156,848 women in the multivariable regression and one-sample Mendelian randomization analysis in UK Biobank (7,784 with a breast cancer diagnosis) and 122,977 breast cancer cases and 105,974 controls from BCAC in the two-sample Mendelian randomization analysis.

**Exposures:** Self-reported chronotype (morning/evening preference), insomnia symptoms and sleep duration in multivariable regression, and genetic variants robustly associated with these sleep traits.

**Main outcome measures:** Breast cancer (prevalent and incident cases in UK Biobank, prevalent cases only in BCAC).

**Results:** In multivariable regression analysis using data on breast cancer incidence in UK Biobank, morning preference was inversely associated with breast cancer (HR 0.95, 95% CI 0.93, 0.98 per category increase) while there was little evidence for an association with sleep duration and insomnia symptoms. Using 341 single nucleotide polymorphisms (SNPs) associated with chronotype, 91 SNPs associated sleep duration and 57 SNPs associated with insomnia symptoms, one-sample MR analysis in UK Biobank provided some supportive evidence for a protective effect of morning preference on breast cancer risk (HR 0.85, 95% 0.70, 1.03 per category increase) but imprecise estimates for sleep duration and insomnia symptoms. Two-sample MR using data from BCAC supported findings for a protective effect of morning preference (OR 0.88, 95% CI 0.82, 0.93 per category increase) and adverse effect of increased sleep duration (OR 1.19, 95% CI 1.02, 1.39 per hour increase) on breast cancer (both estrogen receptor positive and negative), while there was inconsistent evidence for insomnia symptoms. Results were largely robust to sensitivity analyses accounting for horizontal pleiotropy.

**Conclusions:** We found consistent evidence for a protective effect of morning preference and suggestive evidence for an adverse effect of sleep duration on breast cancer risk.

## Introduction

In 2007, the World Health Organization’s International Agency for Research on Cancer (IARC) classified shift work that involves circadian disruption as a probable carcinogen (1). Disturbed sleep, light exposure at night and exposure to other lifestyle factors have been proposed as possible underlying mechanisms (2–4). However, much of the evidence for the carcinogenic effect of shift work comes from animal models rather than epidemiological or experimental studies in humans (1, 4). Furthermore, while the literature on breast cancer risk has focused on the potentially adverse effects of night shift work and exposure to light-at-night, there has been much less investigation into the potential adverse effects of sleep disruption and traits such as chronotype (morning/evening preference), sleep duration and insomnia (5).

In a meta-analysis of 28 studies, there was strong evidence for a positive association between circadian disruption and breast cancer risk (RR 1.14, 95% CI 1.09-1.21). However, the association with short sleep duration (<7 hours per night) in 7 contributing studies was much less consistent (RR 0.96, 95% CI 0.86-1.06) and no dose-response association with sleep deficiency was observed (6). Findings from other meta-analyses have been conflicting, with two showing no consistent evidence that sleep duration is associated with breast cancer risk (7, 8) and one showing evidence of an adverse effect of increased sleep duration (>7 hours) (9). However, the majority of studies in the meta-analyses have tended to be case-control designs, vulnerable to reverse causation, or cohort studies with a small number of cases. Fewer studies have been conducted to investigate associations between chronotype and insomnia with breast cancer risk. A cohort study of 72,517 women (1,834 breast cancer cases) in the Nurses’ Health Study found no strong evidence of association with chronotype (10) and a prospective study of 33,332 women (862 incident breast cancer cases) in the HUNT study found no strong evidence of association with individual insomnia symptoms, although there was evidence of some excess risk among individuals with multiple insomnia complaints (11). Studies have tended to rely on self-report of sleep exposures, meaning associations may be biased by measurement error, and residual or unmeasured confounding in these observational studies makes causal inference challenging.

Mendelian randomization (MR) uses genetic variants that are robustly associated with potentially modifiable risk factors to explore their causal effects (12–14). This method is less susceptible to measurement error, confounding and reverse causation than conventional multivariable regression approaches, provided certain assumptions are not violated. These are that the genetic variants: are strongly associated with the exposure of interest; are not associated with confounders of the exposure-outcome relationship; do not influence the outcome via pathways other than the exposure of interest. Genetic variants robustly associated with chronotype, sleep duration and insomnia symptoms have recently been identified in large genome wide association studies (GWAS) with sample sizes of ^~^50,000 to >1 million participants (15–23). Findings from those GWAS have confirmed the role of several core circadian genes influencing sleep traits and identified genetic variants with no previously known circadian role (24). These genetic variants have been used in two-sample MR and provided some evidence that longer sleep has a causal effect on schizophrenia risk (16), while being a morning person is causally associated with reduced risk of schizophrenia and depression (15), and insomnia is causally associated with increased risk of type 2 diabetes, coronary heart disease and several psychiatric traits (17, 23). To our knowledge MR has not been used to date to explore the causal effect of sleep traits on breast cancer risk.

Using genetic variants robustly associated with chronotype, sleep duration and insomnia symptoms identified in three recent UK Biobank GWAS (15–17), we investigated whether these sleep traits have a causal effect on breast cancer risk. To do this, we performed a one-sample MR analysis using data from UK Biobank, for which estimates were compared with conventional observational multivariable regression results in the same study, as well as a two-sample MR analysis using data from the Breast Cancer Association Consortium (BCAC) (25). Furthermore, we aimed to assess the extent to which findings were robust to potential pleiotropy and supported by genetic variants associated with accelerometer-derived measures of chronotype (sleep-midpoint timing of the least active 5 hours of the day), sleep duration and sleep fragmentation (number of nocturnal sleep episodes).

## Methods

### Multivariable regression and one sample Mendelian randomization analysis

#### Study participants

We used data on women from the UK Biobank, which recruited 502,547 individuals (273,407 women) out of 9.2 million eligible individuals aged between 40 and 70 years in the UK who were invited to participate (5.5% response rate) (26). The study protocol is available online (http://www.ukbiobank.ac.uk/wp-content/uploads/2011/11/UK-Biobank-Protocol.pdf) and more details have been published elsewhere (27). The UK Biobank study was approved by the North West Multi-Centre Research Ethics Committee (reference number 06/MRE08/65) and at recruitment all participants gave informed consent to participate and be followed-up. Information on sleep traits (chronotype, sleep duration and insomnia symptoms), breast cancer case status (prevalent and incident cases with up to 9 years of follow-up), relevant confounding factors and genetic variants are available in UK Biobank.

#### Sleep traits

At baseline assessment, conducted in one of 22 UK Biobank Assessment Centres between 2006 and 2010, participants were given a touchscreen questionnaire, which included questions about sociodemographic status, lifestyle and environment, early life and family history, health and medical history, and psychosocial factors. Participants were asked about their chronotype (morning/evening preference), average sleep duration and any insomnia symptoms.

Chronotype (morning/evening preference) was assessed in the question "Do you consider yourself to be?" with one of six possible answers: “Definitely a ‘morning’ person”, “More a ‘morning’ than ‘evening’ person”, “More an ‘evening’ than a ‘morning’ person”, “Definitely an ‘evening’ person”, “Do not know” or “Prefer not to answer”. A 5-level ordinal variable for chronotype was derived where “Definitely a ‘morning’ person”, “More a ‘morning’ than ‘evening’ person”, “More an ‘evening’ than a ‘morning’ person”, “Definitely an ‘evening’ person”, “Do not know” or “Prefer not to answer” were coded as 2, 1, −1, −2, 0 and missing respectively. Sleep duration was assessed by asking: “About how many hours sleep do you get in every 24 hours? (please include naps)”. The answer could only contain integer values. Binary variables for short sleep duration (<7 hours vs 7-8 hours) and long sleep duration (>8 hours vs 7-8 hours) were also derived. To assess insomnia symptoms, subjects were asked, “Do you have trouble falling asleep at night or do you wake up in the middle of the night?" with responses “never/rarely”, “sometimes”, “usually” or “prefer not to answer”. Subjects who responded “Prefer not to answer” were set to missing. A 3-level ordinal variable for insomnia symptoms was derived where “Never/rarely”, “sometimes” and “usually” were coded as 0, 1 and 2 respectively.

#### Breast cancer

Participants were followed via record linkage to the National Health Service (NHS) Central Registers, which provide information on cancer registrations, coded to the 9^th^ and 10^th^ revision of the International Classification of Diseases (ICD-10), and cancer deaths. The endpoints in these analyses are: i) first diagnosis of invasive breast cancer (ICD-10 C50, ICD-9 174); or ii) breast cancer listed as the underlying cause of death on the death certificate for these women who died during follow-up. We excluded all women with any other cancer diagnosis from the analysis. At the time of analysis, mortality data were available up to February 2016 and cancer registry data up to April 2015. Prevalent cases were defined as those diagnosed with breast cancer prior to date of recruitment to the UK Biobank. Incident cases were defined as those diagnosed with breast cancer or dying from it during follow up.

#### Confounders

We considered the following to be potential confounders of the association between sleep traits on breast cancer risk: education, body mass index, alcohol intake, smoking, strenuous physical activity, family history of breast cancer, age at menarche, parity, use of oral contraceptives, menopause status and menopausal hormone therapy.

BMI was derived from weight and height measured when participants attended the initial assessment centre while information on other potential confounders was obtained from questionnaire responses completed at baseline (**see Supplementary Methods**). Additional information extracted from the initial assessment visit included: centre of initial assessment visit, age at recruitment derived from date of birth and date of attending assessment centre. Participants who were employed were also asked whether their current job involved night shifts never/rarely, sometimes, usually or always.

#### Genetic variants

The full data release in UK Biobank contains the cohort of successfully genotyped individuals (N = 488,377). A total of 49,979 individuals were genotyped using the UK BiLEVE array and 438,398 using the UK Biobank axiom array. Pre-imputation quality control, phasing and imputation of the UK Biobank genetic data have been described elsewhere (28).

In the main analysis, a total of 341 single-nucleotide polymorphisms (SNPs) associated with chronotype (15), 91 SNPs associated with continuous sleep duration (16) and 57 SNPs associated with insomnia symptoms (17), were used in our Mendelian Randomization analysis (**Supplementary Tables 1-3**).

#### Multivariable regression analysis

We carried out multivariable Cox regression between chronotype, insomnia symptoms and sleep duration and incident breast cancer in order to investigate prospective associations between these sleep traits and to minimise the likelihood of reverse causality in observational associations. To minimise the role of confounding, analyses were conducted with adjustment for age, assessment centre and the top 40 genetic principal components (to account for geographical variation). A second model additionally adjusted for degree status, body mass index, alcohol intake, smoking, strenuous physical activity, family history of breast cancer, age at menarche, parity, menopause status, use of oral contraceptives and menopausal hormone therapy.

#### One-sample Mendelian randomization analysis

For one-sample MR, the genetic variants used were extracted genotypes from the UK Biobank Imputation dataset (using genetic variants imputed to the Haplotype Reference Consortium (HRC) reference panel), which had extensive quality control performed including exclusion of the majority of third degree or closer relatives from a genetic kinship analysis (29) (**see Supplementary Methods**). Unweighted allele scores were generated as the total number of sleep trait-increasing alleles (morning preference alleles from chronotype) present in the genotype of each individual.

A two-stage method was implemented where the first stage model consisted of a regression of the sleep trait (chronotype, sleep duration and insomnia symptoms) on the allele score and the second-stage model consisted of a cox regression of breast cancer status on the fitted values from the first-stage regression, to give a population-averaged causal hazard ratio, with adjustment for age, assessment centre, 40 genetic principal components (PCs) and genotyping chip in both stages.

#### Sensitivity analyses

We also performed MR analysis using only those genetic variants which replicated at Bonferroni significance in a large independent dataset for chronotype (15) (242 variants in 23andMe, n=240,098, highlighted in **Supplementary Table 1**) in order to evaluate the potential impact of winner’s curse in the MR analysis. Given the relatively small sample size of replication datasets for sleep duration (CHARGE Consortium, n=47,180) (16) and insomnia (HUNT, n=62,533) (17), few SNPs independently replicated at Bonferroni significance to serve as sufficiently strong instruments for this sensitivity analysis.

To test the MR assumption that genetic variants should not be associated with confounders of the exposure-outcome relationship, we investigated associations between the allele scores and potential confounders in UK Biobank, and performed one-sample MR analysis adjusting for any potential confounders found to be strongly associated with the allele scores as a further sensitivity analysis.

We also conducted both multivariable regression and one-sample MR using all breast cancer cases (incident and prevalent cases) in a logistic regression analysis in UK Biobank and performed sensitivity analysis removing individuals who reported currently working night shifts (sometimes, usually or always).

#### Two-sample Mendelian randomization analysis

We conducted a two-sample Mendelian randomization analysis of sleep traits using: i) female-specific estimates of the associations between the genetic instruments identified in the respective GWAS (15–17) in relation to chronotype (5-level ordinal variable), sleep duration (continuous variable) and insomnia symptoms (3-level ordinal variable) in UK Biobank (**Supplementary Tables 1-3**); and ii) estimates of the associations between the genetic instruments and breast cancer from a large-scale GWAS of breast cancer.

GWAS of chronotype, sleep duration and insomnia symptoms were performed among females on European ancestry (n= 241,350 - 245,767) in the UK Biobank using BOLT-LMM (30) linear mixed models and an additive genetic model adjusted for age, sex, 10 principal components of ancestry, genotyping array and genetic correlation matrix, as was done previously (15–17).

The GWAS of breast cancer involved 122,977 women with breast cancer (estrogen receptor positive (ER+) and estrogen receptor negative (ER-) combined) and 105,974 controls of European ancestry from the Breast Cancer Association Consortium, BCAC (25). BCAC summary data were based on imputation to the 1000 Genomes Project Phase 3 reference panel. To explore potential heterogeneity by breast cancer subtype, we also investigated the causal effect of the sleep traits on breast cancer stratified by estrogen receptor status, using genetic association data from 69,501 ER+ and 21,468 ER- cases within BCAC (25).

Two-sample MR analyses were conducted using MR Base, an R package for two-sample MR (31). We first used an Inverse-variance weighted (IVW) method to meta-analysis the SNP-specific Wald estimates (SNP-outcome estimate ÷ SNP-exposure estimate) using random-effects. If a SNP was unavailable in the breast cancer GWAS summary statistics, then proxy SNPs were identified with a minimum linkage disequilibrium (LD) r^2^ = 0.8. Palindromic SNPs were aligned if they had a minor allele frequency <0.3, or were otherwise excluded.

#### Sensitivity analyses

The IVW random effects method will return an unbiased estimate in the absence of horizontal pleiotropy or when horizontal pleiotropy is balanced (32). To account for directional pleiotropy, we compared results with three other MR methods which each make makes different assumptions about this: MR Egger (33), weighted median (34) and weighted mode (35), and therefore a consistent effect across multiple methods strengthens causal evidence.

To further detect and correct obtained causal estimates for potential violation of the MR assumptions (32), we performed RadialMR (36) in the two-sample context to identify outliers which have the most weight in the MR analysis and the largest contribution to Cochran’s Q statistic for heterogeneity, which may then be removed and the data re-analysed. Radial MR analysis was conducted using modified second order weights and an alpha level of 0.05 ÷ the number of SNPs being used to instrument the exposure.

In order to evaluate the potential impact of winner’s curse, we performed two-sample MR analysis using 242 genetic variants which replicated at Bonferroni significance in a large independent dataset for chronotype (15) (23andMe, n=240,098, highlighted in **Supplementary Table 1**).

Given potential non-linear associations between sleep duration and breast cancer risk (9), we also used data on 27 SNPs associated with short sleep (<7 hours vs 7-8 hours) and 8 SNPs associated with long sleep (>8 hours vs 7-8 hours) (16) in two-sample MR analysis (**Supplementary Tables 4-5**). Causal effect estimates (i.e. ORs for breast cancer) were rescaled to be interpreted per doubling of genetic liability for short or long sleep, as recommended by Burgess et al (37).

Finally, we performed two-sample MR using genetic variants robustly associated with accelerometer-derived sleep traits in UK Biobank, to be compared with causal estimates obtained using genetic variants associated with self-reported traits. For this, we used genetic variants identified in GWAS in relation to three accelerometer based measures: timing of the least active 5 hours (L5 timing) (6 SNPs), nocturnal sleep duration (11 SNPs) and number of nocturnal sleep episodes (21 SNPs) in up to 85,205 individuals, as previously described (38) (**Supplementary Tables 6-8**). More details about how accelerometer sleep traits were derived can be found in the **Supplementary Methods**. Effect estimates represented an hour earlier L5 timing (correlated positively with and to be compared with the self-reported chronotype measure of increased morning preference), an hour increase of nocturnal sleep duration (to be compared with self-reported sleep duration), and a unit increase in the number of nocturnal sleep episodes (to be compared with self-reported insomnia symptoms).

All analyses were conducted using Stata (version 15) and R (version 3.4.1).

## Results

### Baseline characteristics

Of the 180,216 women in the UK Biobank who had been successfully genotyped and passed the genetic QC, and after excluding those who had been diagnosed with other types of cancer, 7,784 (4.9%) had been exclusively diagnosed with breast cancer. Of these, 5,036 (3.2%) were defined as prevalent cases and 2,740 (1.7%) developed incident breast cancer over a median follow-up time of 2.98 years.

Women with a breast cancer diagnosis (prevalent or incident) were more likely to: be older, have a high body mass index, be less physically active, have had an earlier age at menarche, have gone through the menopause, have ever used hormone replacement therapy, to have a family history of breast cancer and be nulliparous; and were less likely to: be never smokers, do night shift work and to have ever used oral contraceptives (**Table 1**), compared with women without a breast cancer diagnosis. There was no strong difference in education level between individuals with and without breast cancer, in line with previous findings (39), as well as no clear difference in drinking status.

**Table 1.** Baseline characteristics of women who had and had not developed breast cancer by date of censoring in UK Biobank.

| Characteristic | Women without breast cancer diagnosis (total n=149,064) | Women with prevalent breast cancer diagnosis (total n=7,784) |
| --- | --- | --- |
| **Age at recruitment (years) (n, mean (SD))** | 149,064<br>56.2 (7.9) | 7,784<br>58.9 (7.0) |
| **Body mass index (n, mean (SD))** | 148,617<br>27.0 (5.1) | 7,764<br>27.2 (4.9) |
| **Age at menarche (n, mean (SD))** | 144,845<br>13.0 (1.6) | 7,576<br>12.8 (1.6) |
| **Days/week strenuous physical activity (n, mean (SD))** | 141,387<br>1.7 (1.8) | 7,325<br>1.5 (1.8) |
| **Education** |  |  |
| Degree (n (%)) | 65,381 (44.3) | 3,337 (43.3) |
| No degree (n (%)) | 89,383 (55.8) | 4,376 (56.7) |
| **Smoking** |  |  |
| Never (n (%)) | 90,072 (60.6) | 4,445 (57.4) |
| Former (n (%)) | 46,374 (31.2) | 2,700 (34.8) |
| Current (n (%)) | 12,131 (8.2) | 604 (7.8) |
| **Alcohol** |  |  |
| Never (n (%)) | 6,254 (4.2) | 343 (4.4) |
| Former (n (%)) | 5,208 (3.5) | 287 (3.7) |
| Current (n (%)) | 137,465 (92.3) | 7,145 (91.9) |
| **Family history of breast cancer** |  |  |
| Yes (n (%)) | 9,221 (6.2) | 526 (6.8) |
| No (n (%)) | 139,843 (93.8) | 7,258 (93.2) |
| **Parity** |  |  |
| 0 (n (%)) | 27,508 (18.5) | 1,489 (19.1) |
| 1+ (n (%)) | 121,479 (82.5) | 6290 (80.9) |
| **Oral contraceptive use** |  |  |
| Yes (n (%)) | 123,688 (83.1) | 6,193 (79.7) |
| No (n (%)) | 25,117 (16.9) | 1,580 (20.3) |
| **Menopause** |  |  |
| Yes (n (%)) | 89,397 (60.0) | 5,721 (73.6) |
| No (n (%)) | 36,272 (24.4) | 787 (10.1) |
| Not sure (n (%)) | 23,277 (15.6) | 1,266 (16.3) |
| **Hormone replacement therapy** |  |  |
| Yes (n (%)) | 57,038 (38.4) | 3,142 (40.5) |
| No (n (%)) | 91,682 (61.7) | 4,620 (59.5) |
| **Shift work** |  |  |
| Night shift (n (%)) | 4,815 (5.8) | 158 (4.6) |
| Other shift (n (%)) | 6,763 (8.1) | 282 (8.2) |
| No (n (%)) | 71,790 (86.1) | 3,014 (87.3) |

### Multivariable analysis

In multivariable Cox regression analysis, we observed an inverse association between morning preference and risk of breast cancer which remained similar in the fully adjusted model (HR 0.95, 95% CI 0.93, 0.98 per category increase in morning preference) but no clear association between sleep duration and insomnia symptoms with risk of breast cancer (**Table 2**). When incident and prevalent cases were combined and associations investigated in a logistic regression framework, there was consistent evidence for an inverse association between morning preference and risk of breast cancer (OR 0.96, 95% CI 0.94, 0.98), as well a positive association between both sleep duration (OR 1.02, 95% CI 1.00, 1.05 per hour increase) and insomnia symptoms (OR 1.11, 95% CI 1.07, 1.15 per category increase) with risk of breast cancer, potentially reflecting reverse causation (**Supplementary Table 9**). Cox regression estimates were similar when individuals who reported working night shifts were excluded (**Supplementary Table 10**).

**Table 2.** Multivariable and Mendelian randomization Cox regression analysis of the risk of breast cancer associated with sleep traits.

| Sleep trait | Basic model* |  |  |  | Fully adjusted model † |  |  |  | Mendelian randomization analysis** |  |  |  |
| --- | --- | --- | --- | --- | --- | --- | --- | --- | --- | --- | --- | --- |
|  | N (cases) | HR | 95% CI | P | N (cases) | HR | 95% CI | P | N (cases) | HR | 95% CI | P |
| Chronotype (per category increase from definite evening, intermediate evening, don't know, intermediate morning, definite morning) | 151,421 (2,732) | 0.94 | 0.92, 0.97 | 7.8x10<sup>-5</sup> | 138,529 (2,500) | 0.95 | 0.93, 0.98 | 0.002 | 151,421 (2,732) | 0.85 | 0.70, 1.03 | 0.098 |
| Sleep duration (per hour increase) | 150,845 (2,723) | 1.01 | 0.98, 1.05 | 0.553 | 138,228 (2,495) | 1.00 | 0.96, 1.04 | 0.984 | 150,845 (2,723) | 1.06 | 0.70, 1.59 | 0.784 |
| Insomnia symptoms (per category increase from none, some, frequent) | 149,005 (2,740) | 1.02 | 0.97, 1.08 | 0.444 | 138,771 (2,505) | 1.02 | 0.97, 1.08 | 0.442 | 149,005 (2,740) | 1.37 | 0.59, 3.20 | 0.468 |

\*Adjusted for age, assessment centre and the top 40 genetic PCs
†Adjusted for age, assessment centre, top 40 genetic PCs, degree status, body mass index, alcohol intake, smoking, strenuous physical activity, family history of breast cancer, parity, age at menarche, menopause status, use of oral contraceptives and menopausal hormone therapy
\*\*Adjusted for age, assessment centre, top 40 genetic PCs and genotyping chip

### One-sample Mendelian randomization analysis

Among UK Biobank female participants, allele scores explained 2.3% of the variance in chronotype, 0.7% of the variance in sleep duration and 0.4% of the variance in insomnia symptoms, respectively (**Table 3**). There was moderate evidence for a protective effect of morning preference on breast cancer risk (HR 0.85, 95% 0.70, 1.03 per category increase) and weaker evidence for an adverse effect of increased sleep duration (HR 1.06, 95% CI 0.70, 1.59 per hour increase) and insomnia symptoms (HR 1.37, 95% 0.59, 3.20 per category increase) (**Table 2**), albeit imprecisely estimated (wide confidence intervals). When using only the genetic variants which replicated in an independent dataset (242 variants in 23andMe) for chronotype, effect estimates of association with breast cancer risk were similar (HR 0.89, 95% CI 0.71, 1.12 per category increase) although with wider confidence intervals given that the replicated variants explained less of the variance in chronotype (1.6%) (**Supplementary Table 11**).

**Table 3.** Allele scores for sleep traits in UK Biobank.

| Sleep trait | N | Mean number of increasing alleles | SD | Association of allele score with sleep trait* |  |  |  |
| --- | --- | --- | --- | --- | --- | --- | --- |
|  |  |  |  | Coefficient (SE) | P-value | R<sup>2</sup> | F-statistic |
| Chronotype (morningness) | 156,454 | 336 | 11.6 | 0.017 (0.0003) | <1x10<sup>100</sup> | 0.0229 | 3666 |
| Sleep duration | 150,845 (2,723) | 90 | 5.9 | 0.015 (0.0004) | <1x10<sup>100</sup> | 0.0072 | 1127 |
| Insomnia symptoms | 149,005 (2,740) | 56 | 4.9 | 0.012 (0.0002) | <1x10<sup>100</sup> | 0.0041 | 639 |

\*Adjusted for age, assessment centre, 40 PCs and genotype chip

While the majority of confounding factors were not associated with the sleep trait allele scores in UK Biobank, after accounting for multiple testing, we found that the chronotype allele score was associated with parity and vigorous activity; the sleep duration allele score was associated with age at menarche and body mass index; and the insomnia allele score was associated with taking hormone-replacement therapy and age at menarche (**Supplementary Table 12**). Further sensitivity analysis was undertaken adjusting for these potential confounders in the one-sample MR analysis and effect estimates were consistent (**Supplementary Table 13**).

Findings of a protective effect of morning preference were supported in analysis of all breast cancer cases (incident and prevalent) in logistic regression. There was weaker evidence for sleep duration and insomnia symptoms, although both had effect estimates in the positive direction (**Supplementary Table 9**). In analyses removing individuals who reported working night shifts, findings were also consistent with the main results from Cox regression (**Supplementary Table 10**).

### Two-sample Mendelian randomization analysis

After harmonization of the SNP effects in the two summary datasets (UKBiobank and BCAC), 305 SNPs were used to instrument chronotype, 82 SNPs for sleep duration and 50 SNPs for insomnia symptoms. Findings of a protective effect of morning preference were supported by two sample MR (IVW OR 0.88, 95% CI 0.82, 0.93 per category increase) (**Supplementary Table 14, Supplementary Figure 1**) as well as an increased risk of sleep duration (IVW OR 1.19, 95% CI 1.02, 1.39 per hour increase) (**Supplementary Table 14, Supplementary Figure 2**). Little evidence for a causal effect of insomnia symptoms was observed (IVW OR 0.93, 95% CI 0.49, 1.76 per category increase) (**Supplementary Table 14, Supplementary Figure 3**). IVW estimates for chronotype, sleep duration and insomnia symptoms from two-sample MR are compared with multivariable and one-sample MR approaches in UK Biobank in **Figure 1**. Findings were similar when stratified by ER+ and ER- breast cancer (**Supplementary Table 14**).

**Figure 1.**
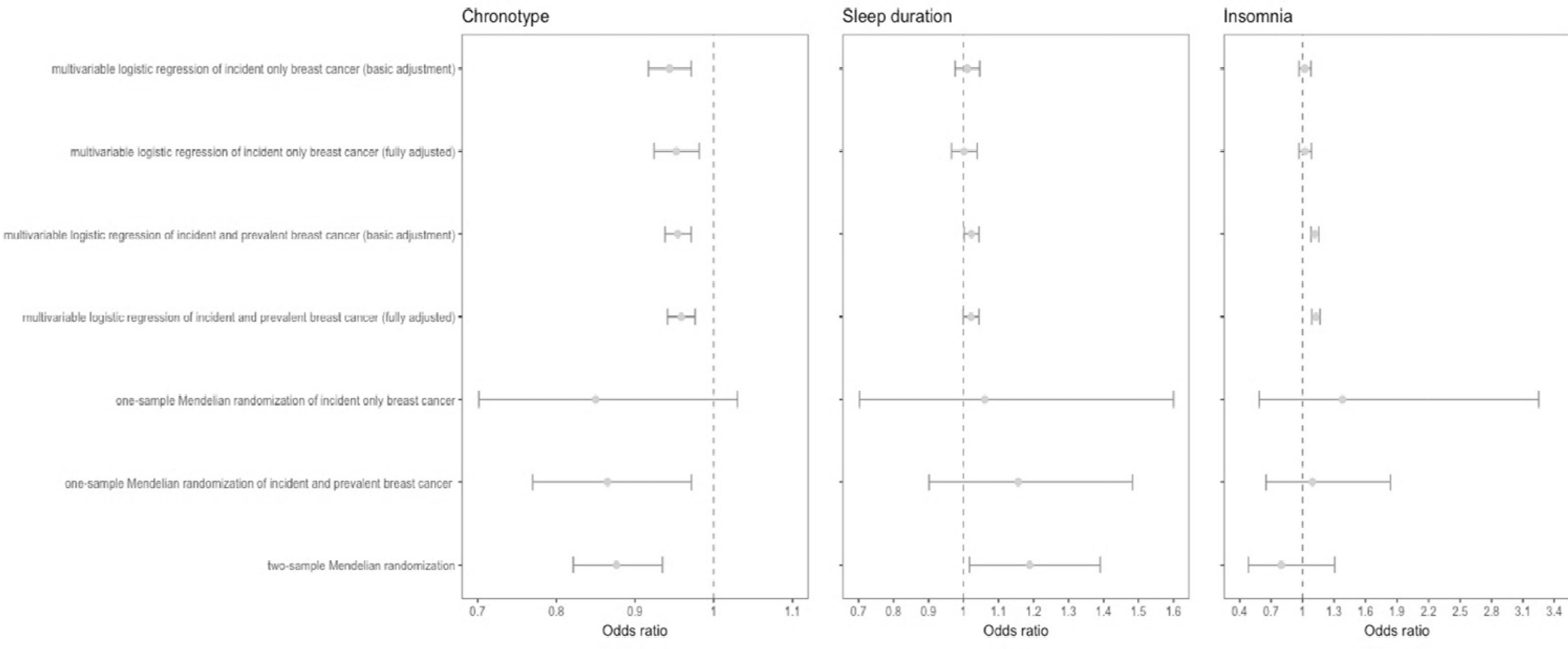
Forest plot of Multivariable and Mendelian randomization estimates for the association between sleep traits and breast cancer risk. Odds ratios are per category increase in chronotype (from (i) definite evening (ii) intermediate evening (iii) neither (iv) intermediate (v) definite morning), per hour increase in sleep duration and per category increase in insomnia risk (from (i) no insomnia symptoms insomnia symptoms (iii) frequent insomnia symptoms). N.B. odds ratios rather than hazard ratios for incident breast cancer are show multivariable and one-sample Mendelian randomization analysis in order to compare estimates across methods.

Effect estimates were broadly consistent between the IVW method and the pleiotropy-robust methods applied (MR-Egger, Weighted Median and Weighted Mode) in two-sample MR (**Supplementary Table 14, Figures 1-3**). Furthermore, the MR Egger test of directional pleiotropy was consistent with the null for all analyses (**Supplementary Table 15**).

Evidence for heterogeneity in causal effects for most of the models (**Supplementary Table 16**) could still indicate potential violations of the MR assumptions. We used radial plots to aid in the detection of outlying variants. Radial MR analysis identified 6 outliers for chronotype, 3 outliers for sleep duration and 2 outliers for insomnia symptoms in both IVW and MR Egger (**Supplementary Table 17, Supplementary Figures 4-6**). With outlier removal, IVW and MR Egger effect estimates were largely unchanged (**Supplementary Table 18**).

Effect estimates for the causal effect of chronotype on breast cancer were consistent when using the 242 genetic variants associated with chronotype which replicated at Bonferroni significance in 23andMe (15), indicating that winner’s curse is unlikely to have substantially biased effect estimates (**Supplementary Table 19**).

Findings of an adverse effect of increased sleep duration on breast cancer risk were supported using genetic variants specifically associated with short and long sleep duration, with evidence for a protective effect of short sleep duration on breast cancer (IVW OR 0.92, 95% CI 0.86, 0.99 per doubling of genetic liability for short sleep duration) and adverse effect of long sleep duration (IVW OR 1.24, 95% CI 0.96, 1.60 per doubling of genetic liability for long sleep duration) (**Supplementary Table 20**).

Using genetic variants robustly associated with accelerometer-derived sleep traits in UK Biobank, we found no clear evidence of association with L5 timing measured objectively (IVW OR 1.04, 95% CI 0.78, 1.38 per hour decrease) (**Supplementary Table 21, Supplementary Figure 9**). However, an adverse effect of increased sleep duration was supported using estimates from objectively-measured sleep duration (IVW OR 1.16; 95% 1.02, 1.32 per hour increase) (**Supplementary Table 21, Supplementary Figure 9**) and there was some evidence for a causal effect of increased fragmentation on breast cancer risk (IVW OR 1.14; 95% CI 1.00, 1.30 per sleep episode) (**Supplementary Table 21, Supplementary Figure 10**).

## Discussion

### Overall findings

In this study, we compared observational estimates from multivariable regression to those from Mendelian randomization analyses to make inferences about the likely the causal effects of three sleep traits on breast cancer risk. In multivariable regression analysis using data on breast cancer incidence in the UK Biobank study, morning preference was inversely associated with breast cancer while there was little evidence for an association with sleep duration and insomnia risk. Using genetic variants associated with chronotype, sleep duration and insomnia symptoms, one-sample MR analysis in UK Biobank provided some evidence for a protective effect of morning preference but imprecise estimates for sleep duration and insomnia. Findings for a protective effect of morning preference and adverse effect of increased sleep duration on breast cancer (both estrogen receptor positive and negative) were supported by two-sample MR using data from BCAC, while there was inconsistent evidence for insomnia symptoms. Results were largely robust to sensitivity analyses accounting for horizontal pleiotropy.

This study represents the first Mendelian randomization analysis to investigate the causal effect of sleep traits on risk of breast cancer. One previous study found a strong association between circadian pathway genetic variants and risk of breast cancer (40). Nonetheless, this previous study was unable to directly implicate modifiable sleep traits by which risk of breast cancer could be minimized and did not attempt to separate the effects of the genetic variants on breast cancer risk via circadian disruption from pleiotropic pathways.

Findings of an adverse effect of evening preference on breast cancer risk in all analyses performed are consistent with those studies identifying nightshift work as a potential carcinogen (1) and support hypotheses around carcinogenic light-at-night (4). However, findings when using an objective measure of chronotype (L5 timing) did not reveal the same adverse effect. While the latter analysis may be limited by the number and strength of the genetic variants used to instrument L5 timing, the lack of consistency in estimates draws to question the mechanisms by which morning/evening preference (rather than actual activity) has an effect on breast cancer risk.

Evidence for an adverse effect of increased sleep duration on breast cancer risk is in contrast to observational findings in UK Biobank as well as much of the literature on circadian disruption and breast cancer risk (6). However, recent studies implicate longer sleep duration as a risk factor for breast cancer (9). Given previous reports of a J-shape relationship between sleep duration and breast cancer risk (9), as well as investigating sleep duration as a continuous variable, we also investigated the causal effects of both short and long sleep duration in order to investigate non-linear effects. In line with our main findings, we found evidence for a protective effect of short sleep duration and adverse effect of long sleep duration on breast cancer risk. Furthermore, using genetic variants associated with accelerometer-derived nocturnal sleep duration, we found evidence for an adverse effect of sleep duration with a similar magnitude of effect.

Overall, we found inconsistent evidence regarding the causal effect of insomnia symptoms on breast cancer risk in multivariable and Mendelian randomization analyses. A previous study of incident breast cancer in the HUNT study revealed no strong evidence of association with individual insomnia symptoms (11), although individuals with multiple insomnia complaints were found to be at increased risk. In our analysis, insomnia was defined based on self-report of either difficulty initiating sleep or waking in the night. Therefore further work is required to investigate individual symptoms of insomnia on breast cancer risk, and their potential cumulative effect is required. Interestingly, MR analysis did provide some evidence for adverse causal effect of accelerometer-derived number of nocturnal sleep episodes on breast cancer.

### Strengths and limitation of the study

Key strengths of the study are the integration of multiple approaches to assess the causal effect of sleep traits on breast cancer, the inclusion of data from two large epidemiological resources, UK Biobank and BCAC as well as use of data derived from both self-reported and objectively-assessed measures of sleep. Furthermore, for Mendelian randomization analysis we used the largest number of SNPs identified in the GWAS literature with full summary statistics available in order to obtain strong genetic instruments for MR analysis and to explore potential pleiotropic pathways.

The approaches of multivariable Cox regression of incident cases, multivariable logistic regression of prevalent and incident cases, one-sample Mendelian randomization and two-sample Mendelian randomization, each have different strengths and limitations in terms of key sources of bias (**Table 4**). In multivariable analysis, attempts were made to mitigate key sources of bias, including confounding and reverse causation, with the use of multivariable Cox regression analysis of incident breast cancer cases and adjustment for a number of hypothesised confounders. Nonetheless, residual or unmeasured confounding, selection bias and measurement error may also have distorted effect estimates. We used Mendelian randomization analysis to minimise the likelihood of measurement error, confounding and reverse causation. In addition, we conducted a series of sensitivity analyses to test the core assumptions that the genetic instruments are strongly associated with the exposures of interest, are not influenced by confounding factors and do not directly influence the outcome other than via the exposure.

**Table 4.**
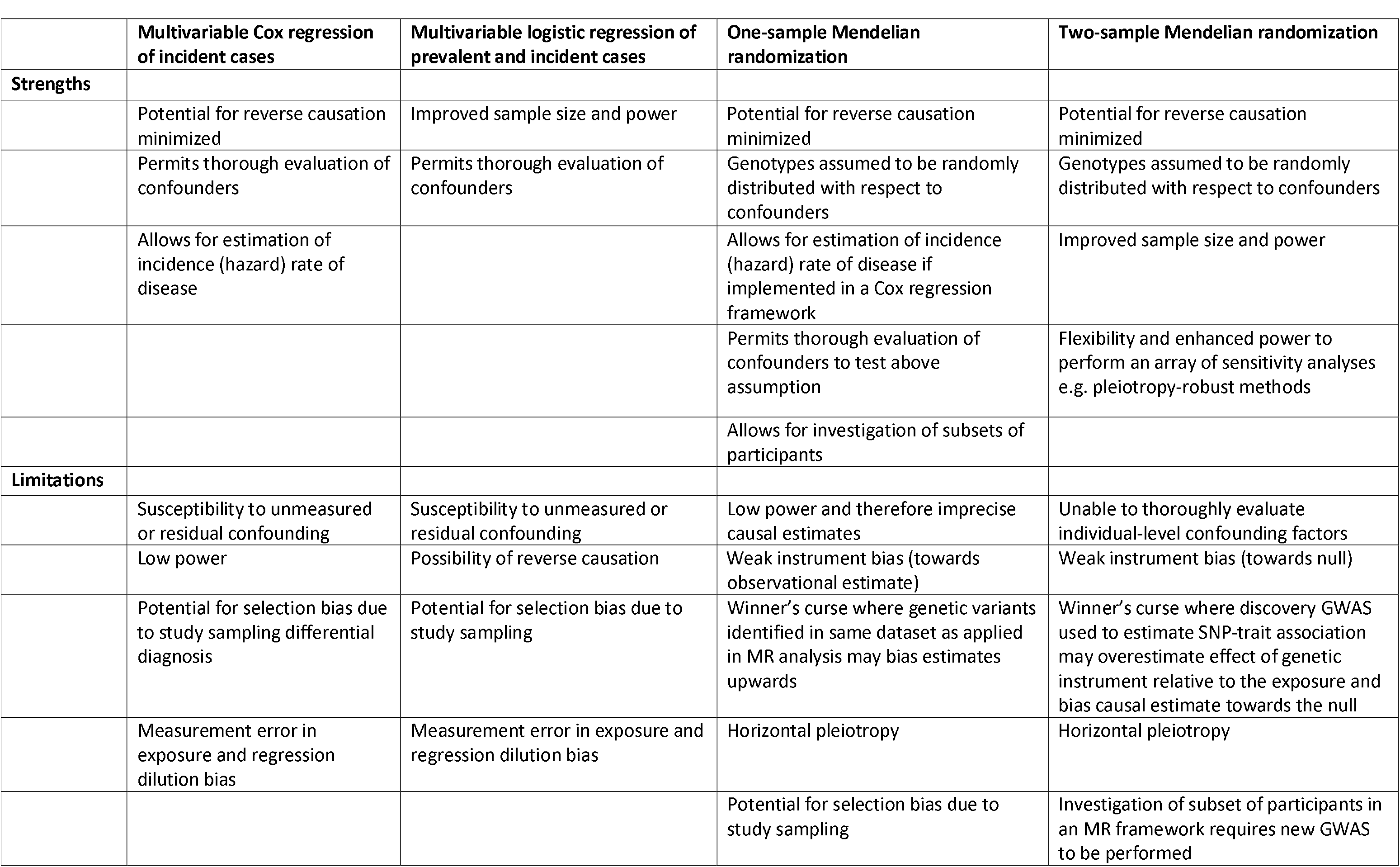
Strengths and limitations of epidemiological approaches applied in this study.

One limitation of this study is related to the selection of participants. Analysis in the two large epidemiological studies included here (UK Biobank and BCAC) was restricted to females of European ancestry. Further work is required to investigate whether these findings translate to individuals in other ancestry groups. While the UK Biobank represents a large and well characterised epidemiological resource, it is not representative of the UK population as a whole given low participation rates (27). As well as influencing the generalisability of findings, selection into the study can lead to biased estimates of association through “collider bias” (41). To minimise the influence of this, we also used genetic data from a large case-control study of breast cancer (BCAC), and compared MR effect estimates across these datasets.

In all MR analyses, SNP-exposure estimates were obtained from the UK Biobank as this has formed a major component of the GWAS of sleep traits conducted to date (15–17, 20, 21, 23). This may lead to ‘winner’s curse’ where the magnitude of the effect sizes for genetic variants identified within a discovery sample are likely to be larger than in the overall population. In a one-sample MR analysis, the impact of winner’s curse of the SNP-exposure association can bias causal estimates towards the confounded observational estimate while in two-sample MR, winner’s curse can result in bias of the causal estimate towards the null. To minimise the impact of winner’s curse in one-sample MR analysis we derived an additional allele score for chronotype composed of SNPs which replicated beyond a Bonferroni-correction threshold in an independent study (23andMe) (15). Similarly, for two-sample MR analysis, we used SNP-exposure estimates from this replication analysis in sensitivity analyses and findings were consistent with the main analysis (**Supplementary Table 11 and 19**).

We were unable to apply the same approach to investigate the impact of winner’s curse in the sleep duration and insomnia analysis due to the relatively small sample size of the replication datasets in those studies, meaning genetic associations could be imprecise. While we are aware of a large GWAS for insomnia which was conducted using data from both UK Biobank and 23andMe, full summary data for the top SNPs identified in this analysis have yet to be published (23). We used unweighted allele scores to minimise the contribution of potential weak instruments in the one-sample Mendelian randomization analysis.

While associations between the allele scores and confounders in UK Biobank imply violation of the MR assumption that genetic variants should not be associated with confounding factors, there are several explanations for these findings. Previous MR studies have identified causal effects of sleep traits on reproductive traits and activity levels (15–17) suggesting that these factors may be mediators of the association between sleep traits and breast cancer rather than confounders. Furthermore, some of the genetic variants associated with chronotype and insomnia have been found to be adiposity-related loci (15, 16), implying potential pleiotropic pathways. Nonetheless, we also applied a series of pleiotropy-robust MR methods and outlier detection to rigorously explore the possibility that findings of a causal effect of chronotype and sleep duration were not biased due to pleiotropy.

As well as attempting to mitigate key sources of bias for each epidemiological approach applied, we have also assessed the consistency in estimates between the approaches in order to provide the best inference regarding the causal effect of sleep traits on breast cancer. This is aligned with the practice of “triangulation” which aims to obtain more reliable answers to research questions through the integration of results from different approaches, where each approach has different sources of potential bias that are unrelated to each other (42, 43). We also compared estimates based on self-reported sleep with the use of genetic variants associated with accelerometer-derived measures of sleep (38), although we did not use female-specific SNP estimates here given the smaller number of individuals in UK Biobank with these data.

### Policy implications

Findings of a protective effect of morning preference on breast cancer risk add to other evidence from Mendelian randomization supporting a possible beneficial effect of morning preference on decreased risk of schizophrenia and depression (15). They also support hypotheses around the carcinogenic effect of night shift work and light-at-night (1). However, whether it is the actual behaviour which poses the health risk, or the preference to morning versus eveningness requires further evaluation. While previous attention has been directed at minimizing the impact of night shift work, here we found some evidence for a detrimental effect of evening preference even among non-shift workers, pointing to other types of public health interventions targeted at the general population. Furthermore, suggestive evidence for a causal effect of increased sleep duration on breast cancer risk should also be taken into account when designing interventions which influence sleep habits of the general population in order to improve health.

### Conclusions

In this study, both multivariable regression and Mendelian randomization analysis were used to provide strong evidence for causal effect of chronotype on breast cancer risk. Furthermore, there was some evidence for a causal effect of both sleep duration on risk of breast cancer, although findings for these traits were less consistent across the different methods applied. However, the biological role of many of the genetic variants used to instrument these traits in Mendelian randomization and mechanistic pathways underlying the observed effects are not well understood. Previously reported pathways between sleep disruption and mammary oncogenesis include immunological, molecular, cellular, neuroendocrine and metabolic processes (5). Further work to uncover these possible mediating processes is required. Nonetheless, these findings have potential policy implications for influencing sleep habits of the general population in order to improve health.

## Funding

The breast cancer genome-wide association analyses were supported by the Government of Canada through Genome Canada and the Canadian Institutes of Health Research, the ‘Ministère de l’Économie, de la Science et de l’Innovation du Québec’ through Genome Québec and grant PSR-SIIRI-701, The National Institutes of Health (U19 CA148065, X01HG007492), Cancer Research UK (C1287/A10118, C1287/A16563, C1287/A10710) and The European Union (HEALTH-F2-2009-223175 and H2020 633784 and 634935). All studies and funders are listed in Michailidou et al (2017).

RCR, ELA, JB, CLR, RMM, MM, DAL and GDS are all members of the MRC Integrative Epidemiology Unit at the University of Bristol funded by the MRC (MM_UU_00011/1, MC_UU_00011/2, MC_UU_00011/5, MC_UU_00011/6 and MC_UU_00011/7). RCR is a de Pass VC Research Fellow at the University of Bristol. This study was supported by the NIHR Biomedical Research Centre at the University Hospitals Bristol NHS Foundation Trust and the University of Bristol. The views expressed in this publication are those of the authors and not necessarily those of the NHS, the National Institute for Health Research or the Department of Health and Social Care. This work was also supported by CRUK (grant number C18281/A19169) and the ESRC (grant number ES/N000498/1). SEJ is funded by the Medical Research Council (grant MR/M005070/1). TMF is supported by the European Research Council (grant 323195:GLUCOSEGENES-FP7-IDEAS-ERC). MNW is supported by the Wellcome Trust Institutional Strategic Support Award (WT097835MF).

## Acknowledgements

This research has been conducted using the UK Biobank Resource under Application Numbers 9072, 6818, 15825 and 16391. We would like to thank the participants and researchers from the UK Biobank who contributed or collected data. The authors would like to thank Dr Ruth Mitchell, Dr Gibran Hemani, Mr Tom Dudding and Dr Lavinia Paternoster for conducting the quality control filtering of UK Biobank data and Mr Wes Spiller, Dr Jie Zheng, Dr Gibran Hemani and Dr Philip Haycock for help with data acquisition and statistical analysis. This study has been made possible with the support of Dr Jonathan de Pass and Mrs Georgina de Pass.

